# Collective Almost Synchronization Modeling Used for Motor Imagery EEG Classification

**DOI:** 10.1101/2023.08.23.554551

**Authors:** Thi Mai Phuong Nguyen, Minh Khanh Phan, Yoshikatsu Hayashi, Murilo S. Baptista, Toshiyuki Kondo

**Affiliations:** Department of Computer and Information Sciences Tokyo University of Agriculture and Technology, Tokyo, Japan; Tokyo Teches, Tokyo, Japan; Department of Biomedical Science and Biomedical Engineering, University of Reading, Reading, United Kingdom; Institute for Complex System and Mathematical Biology, University of Aberdeen, Aberdeen, United Kingdom

## Abstract

Classification based on feature extraction is a crucial technique to develop Brain Computer Interface (BCI) systems. The human brain can be considered as a dynamical system, and its behavior measured by EEG signals can be modeled by a group of nonlinear oscillators. Exploring the dynamical nature of EEG signals along with model based approach may improve classification accuracy in BCI. This study proposes a novel feature extraction method for the classification of Motor Imagery (MI) EEG using a dynamical network model operating in a special collective state, so called Collective Almost Synchronization (CAS). The CAS, the nonlinear oscillators set to operate in a weakly coupled regime, can be used to model an EEG. Purpose of this study is to investigate the performance of the CAS model to identify features for the classification of MI states. To achieve this goal, a linear regression method is used and linear coefficients are extracted as feature vectors. Our approach boils down to identifying patterns in the MI-EEG by associating them to the coefficients of a linear regression (or weights of an output function) constructed to model the MI-EEG signals from simulated time-series generated by a dynamical neural network. The dataset 2b from BCI Competition-IV was used to evaluate the performance of the proposed method. Results indicate that the CAS-based classification method is more robust in extracting distinguishable features from EEG signals as compared with other state-of-the-art methods. The proposed method achieved better performance on two-class MI classification. Moreover, the method developed in this study for MI classification across subjects is effective with 74.03% of the accuracy.

## Introduction

EEG is a useful biological signal to distinguish different brain diseases and mental states. It is an technique for recording human brain signals, and is crucial for the Brain-Computer Interface (BCI). In BCI research, Motor Imagery (MI) is an important topic, which reflects the process in which a person imagines performing a certain task, i.e., intention to perform hand or leg movements. Detecting different MI tasks from EEG signals has attracted much attention by researchers, and several feature extraction methods and classifiers were suggested to recognize imagery action [1]. In recent years, there have been several studies aimed to use MI for wheelchair control [2, 3], neuronal game [4], robotic hand control [5, 6], and autonomous driving [7, 8].

Moreover, EEG pattern recognition is significantly more appealing than facial and speech-based recognition methods, given that internal nerve fluctuations cannot be deliberately masked or controlled [9]. In previous studies, subject-independent EEG classification was shown to be difficult to achieve compared with classification from EEG of subject-dependent ones [10]. Therefore, there is great interest to understand how to improve the performance of EEG classification across subjects. Lu et al. proposed to use a deep learning scheme based on a restricted Boltzmann machine to learn EEG features for MI classification [11]. The best recognition accuracy across subjects was 70%, significantly lower than 84% achieved using subject-dependent MI classification. In the work from [12], a supervised learning-based method called HS-CNN reached 87.6% of subject-dependent classification using dataset 2b of BCI Competition-IV [10], but its performance drops as 65.3% in a subject-independent setting.

To overcome this problem, EEG contains intense noise level, high non-stationarity, and non-linearity. All these are great challenges to find effective representations for EEG data features [13]. In MI-EEG signal decoding, several studies have proposed such as spatial feature (common space pattern (CSP) [14]), time-frequency feature (Hilbert-Huang transform [15], discrete wavelet transform [16], empirical mode decomposition [17, 18], etc.), frequency feature (power spectral sensity [19], fast Fourier transform (FFT) [6, 20]), time-frequency image (short-time Fourier transform (STFT) [21], wavelet transform [22, 23]), and linear coefficient [24–26].

A long-range temporal correlation gave a high accuracy for all motor execution and MI classification [27]. More recently, researchers suggested combining the time domain and frequency domain methods to describe the characteristics of MI-EEG [21] more effectively. However, linear methods failed to address the non-linear characteristic of EEG signals [28]. Human brain can be considered as a complex network of connected nonlinear dynamical systems. Recently, it was shown that the brain activity measured by electroencephalogram (EEG) signals could be modeled by nonlinear oscillators [29, 30]. This opens up an idea to propose feature extraction methods that explores the dynamical nature of EEG signals, and which can improve classification accuracy in BCI. Several aspects of the Collective Almost Synchronization (CAS) method have been considered in EEG modelling [30, 31]. CAS is a phenomenon characterized by the existence of a local cluster of neurons possessing roughly constant local mean fields, a consequence of the fact that neurons are very weakly connected [32]. This phenomenon requires computational models of dynamical systems such as Hindmarsh-Rose neurons connected by small coupling strengths to allow the weak interactions between neurons. Similar to what is done in the AR modelling approach, experimental EEG signal can be reproduced by a weighted linear composition of a set of selected neurons in a simulated network of weakly connected HR neurons [31]. The coefficients of the linear composition modelling the experimental EEG signal are then used as the feature vectors in the BCI system [33, 34]. Extracted features are then input to a suitable classifier to perform the final recognition of the state of MI. Our approach to classify is based on a two-step process. First, we extract the features of the output of simulated neural network that reproduces the experimental EEG signal, and then we use these features as the input for standard machine learning-based classifier.

Recently, Deep Learning (DL) received much attention for its superior performance. Convolution Neural Network (CNN) is one of the DL methods, which has shown a remarkable success concerning image classification and computer vision [35]. CNN is also widely used for EEG classification [36, 37]. Several studies showed that it is suitable for complex EEG recognition tasks [38–41]. This classifier can also be applied to various BCI paradigms such as MI and emotion classifications, and obtained competitive high accuracy to state-of-the-art methods [38, 42–44].

To this end, this study proposes a novel EEG feature extraction method using our CAS-based nonlinear network modeling. We use a MI-EEG dataset 2b from BCI competition-IV [45] and set both intra-subject and across-subjects frameworks classification. To the best of our knowledge, this study is the first to show an application of a large dimensional nonlinear dynamical system constructed as a model of the brain to extract features expressed in MI-EEG signals.

The main contributions of the study are summarized as follows:

1. **Feature extraction by CAS**: We propose a novel EEG feature extraction method in which EEG signals are modeled by a complex network of chaotic Hindmarsh-Rose (HR) neurons that are weakly connected and behaving in the CAS state. The weight coefficients of the model are used as an EEG feature.
2. **Classification by CNN**: A CNN is designed to solve the classification problem. The performance of the proposed method is compared with state-of-the-art methods in both intra-subject and across-subjects MI-EEG classification.

The remainder of this paper is organized as follows. In Section 2, we mainly represent the details of the proposed methods, including feature extraction, feature standardization and classifier. In Section 3, dataset and the details of experimental study are introduced. The experimental results and discussion are presented in Section 4. In Section 5, we draw a conclusion for the whole article.

### Proposed scheme

Specific brain activity recorded via EEG has its own frequency band. The MI activities frequency band which is widely used is 8-35 Hz [11, 46]. Our approach is to use the multi-sub-bands decomposition. Regarding Figure 1, the diagram of the proposed method follows steps:

**Fig 1.**
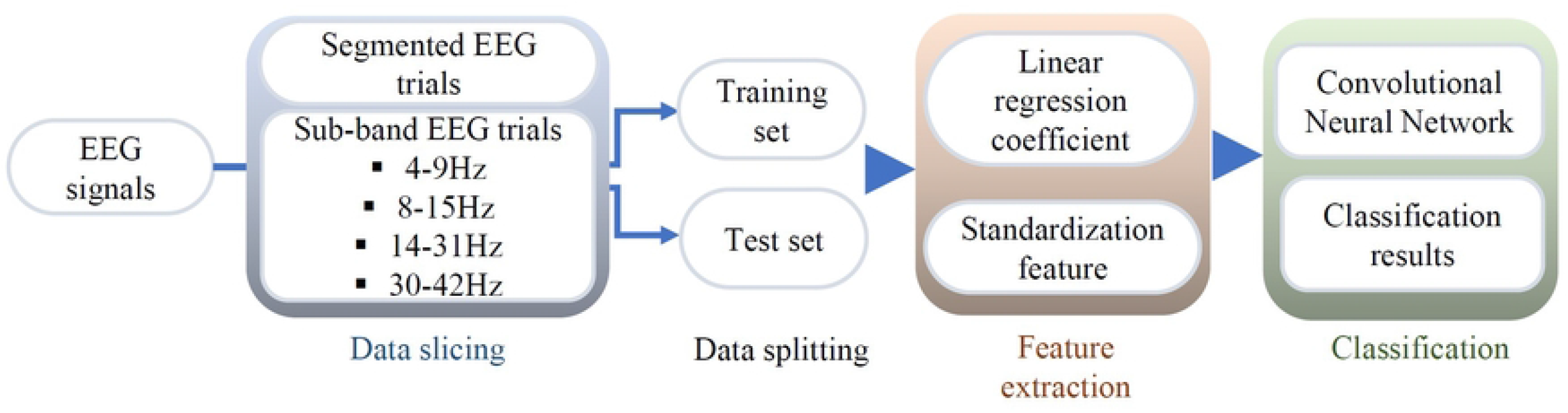
Block diagram of the proposed method.

- Multichannel EEG signal was decomposed into four sub-bands: 4-9 Hz, 8-15 Hz, 14-31 Hz, and 30-42 Hz, then the data was split into training and test data sets.
- CAS-based features were extracted from each sub-band EEG signals.
- The features obtained from multi-sub-bands were combined and standardized. The feature referred to the coefficients to match the EEG signals with states of the neurons in the HR network by a linear regression.
- The CNN classifier was trained with extracted features in the training dataset, and classification test was performed using the test dataset.

### Feature extraction

CAS is presented in complex networks for very weak coupling strength. In CAS, the neuron networks can process infinite possible oscillatory patterns. CAS phenomenon is thus a plausible explanation for the existence of a local cluster of neurons that are sufficiently decorrelated to independently process information locally. Those infinitely many patterns could provide the basis and shed light on the process of memory formation in the brain. Therefore, CAS is defined as a universal way of how patterns can appear in complex networks with nodes connected by small coupling strengths. The HR neuron model is a well-known model for designing the neuron activity. Some studies used HR neuron model to describe a CAS pattern models the EEG signal [30, 31]. In this study, we considered a HR network formed by *L* = 1000 neurons. The HR neurons network model is as below:

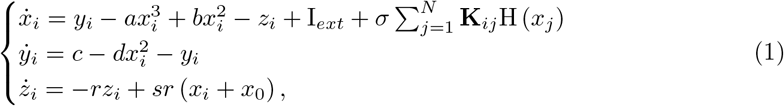

where (*x*_*i*_, *y*_*i*_, *z*_*i*_) ∈ ℝ^3^ are the state variables of the neuron *i, i* = 1 : *N* . *N* is the number of neurons in the network. The parameters of model were chosen as following:*a* = 1, *b* = 3, *c* = 1, *d* = 5, *r* = 0.005, *x*_0_ = 1.618, *I*_*ext*_ = 3.23, and the coupling strength *σ* = 0.001. *K*_*ij*_ was a small-world matrix generated by Watts-Strogatz network approach with a rewiring probability equal to 0.01. **K**_*ij*_ is an adjacency matrix. The HR neuron network with the CAS pattern is described by

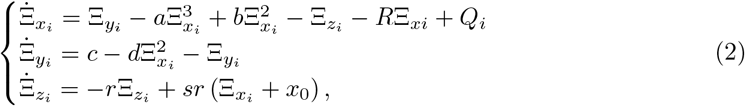

where *R*_*i*_ = *p*_*i*_, *Q*_*i*_ = *p*_*i*_*C*_*i*_, *p*_*i*_ = *σk*_*i*_ and *C*_*i*_*≈* (1*/k*_*i*_) 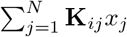.

The HR neurons network with the CAS pattern was simulated using the Brain Dynamics Toolbox consisting *L* = 1000 nodes, and time-series were obtained with the same length of that of the EEG signal time-points, and given by *N* . The network had a mean degree 10 for each node. After that, we record the value of the membrane potential for all *N* neurons and collect *L* data points, from our simulated HR neural network, and store in the matrix *X* =∈ ℝ^*L×N*^ reduced by Principal Component Analysis (PCA) to 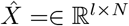 (with *l < L*) and this new reduced set was used to model the EEG signal. Where N is the number of data points. For more details see Refs. [31].

We applied a parametric method using a linear regression model for EEG. Let 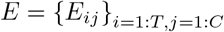, where 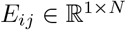, is the sub-band EEG training trials. Here, T is the number of trials, C defines a number of channels. The linear regression model for EEG is defined as below:

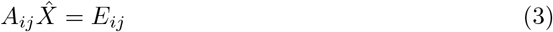

where *A*_*ij*_ = [*α*_0_, *α*_1_, …, *α*_*l*_] is the vector of coefficients, which we regard as the feature extracted from the MI-EEG signals for the trial *i* and for the channel *j*. 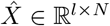 corresponds to PCA reduced state of the HR neuron network operating in CAS regime. The estimation of *A*_*ij*_ is calculated by using:

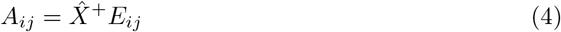

where 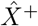 is the Moore-Penrose pseudoinverse of 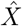. Once *A*_*ij*_ is calculated, an EEG signal can be modelled by

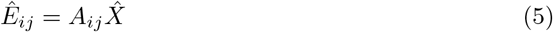

### Feature standardization

Normalization or standardization is a critical step needed to change features to the same scale. Different from normalization, standardization requires the data has a normal distribution. The values of *α*_0_ have changed much for different experiments, and this variation would have resulted in a distribution of the coefficients skewed. Therefore, the out layer *α*_0_ was then removed from *A*_*ij*_. After confirming that the features satisfy the requirement, they were standardized as follows:

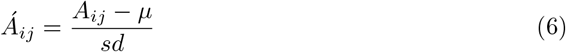

where *A*_*ij*_ is the original feature value, 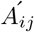 is the normalized feature value. *μ* is the mean and *sd* is the overall standard deviation of the features.

### 0.1 Convolutional Neural Network classifier

CNN can take multidimensional data as input and generally works well for image classification. Various studies used a deep learning approach such as CNN to classify EEG signals. Based on their experimental results, we argue that CNN-based methods significantly improves on classical classification methods [37, 39, 47]. In the present study, we used a simple CNN architecture shown in Figure 2. It had two contiguous blocks of a convolution layer followed by a max-pooling layer, a flatten layer, and finally a dense layer. The CNN model was trained with the SGD optimizer algorithm. Also, we used regularization techniques such as dropout to prevent over fitting problem. The settings used for training the CNN model are as follows:

**Fig 2.**
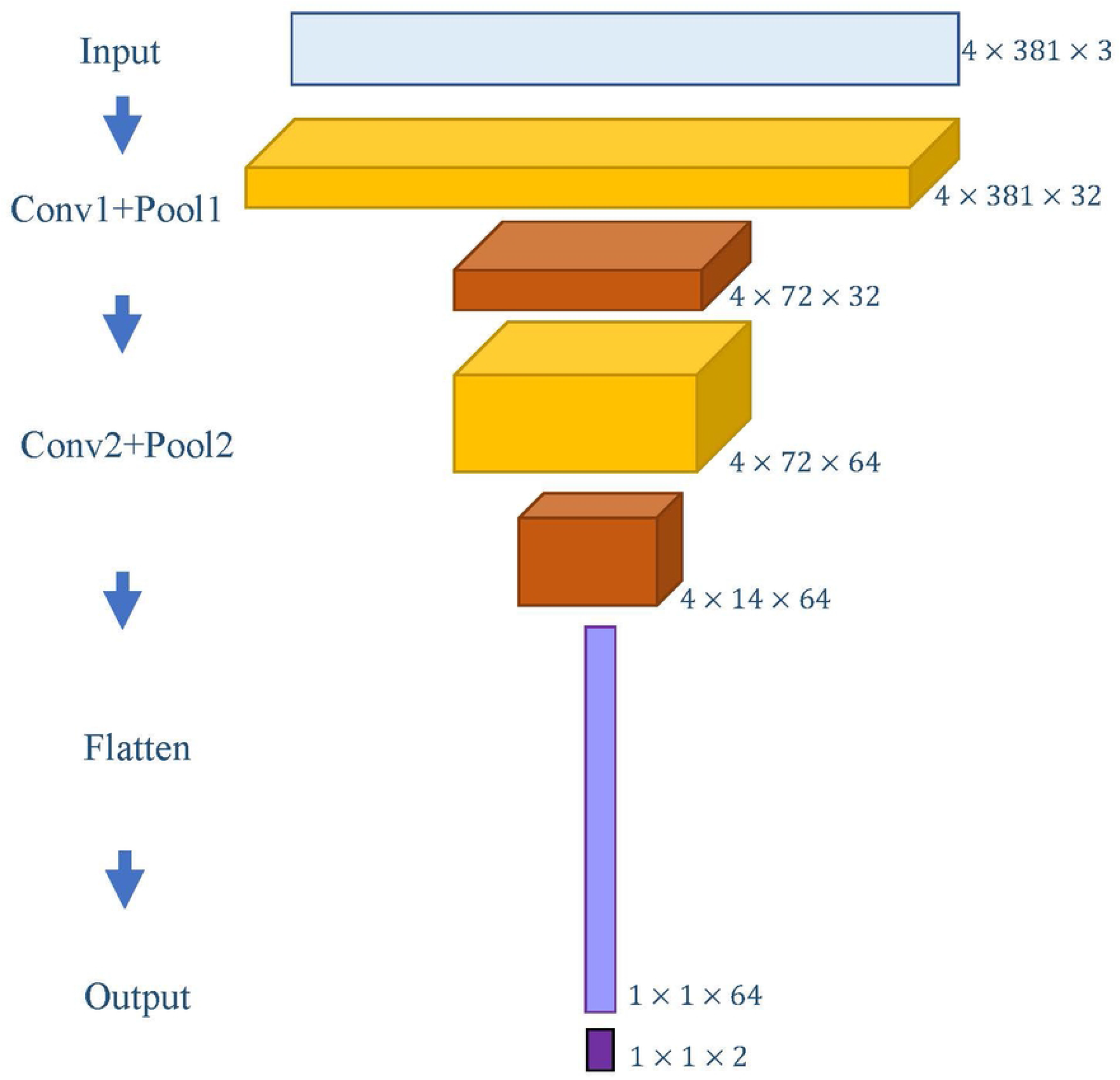
The illustration of the CNN block diagram. The model consists of two convolutional layers, two max pooling layers as well as flatten layer.

- ReLU was used as an activation function in the convolution layer..
- The final fully connected layer derived the probabilities for two output classes using the sigmoid function.
- Stochastic gradient descent with an initial learning rate of 0.01.
- Dropout with a probability of 0.2, 0.3, and 0.4 in each convolution layer and flatten layer, respectively.

## Experiments

### Dataset

To evaluate the performance of the proposed method, we executed MI-EEG classification experiments. For this aim, we used the BCI competition IV dataset 2b [48], because it is commonly used in this field. The dataset contains two types of experiments from nine healthy participants. Each experiment was recorded on separate days for one subject. The MI classification task aims to predict the label of the second experiment from the supervised information of the first experiment.

EEG signals were recorded in three channels: C3, Cz, and C4, with a frequency 250 Hz. Participants executed five recording sessions in total. In the first experiment, two sessions were done without feedback with 120 trials per session, and the rest three sessions in the second experiment incorporated online feedback with 160 trials per session. An EEG time segment was extracted from the first 4 to 7 s, resulting in 750 data points per trial.

In the pre-processing stage, raw EEG data were segmented into trials and band-pass filtered into four sub-bands: theta (4-9Hz), alpha (8-15Hz), beta (14-31Hz), and gamma (30-42Hz). The images achieved by multi-frequency bands representation of EEG signals from a variety of channels are used as the input to the deep neural network. Many studies decomposed EEG signals into bands for classification of MI [10, 11, 49]. The narrow-band signals contains the significant information about movement imagination which can improve the MI classification performance [46].

### Experimental setup and comparative experiments

HR neuron network *X* ∈ ℝ^1000*×*750^ was simulated using Brain Dynamics Toolbox [50]. An example of the dynamic behavior of a single HR neuron is shown in Figure 3. *X* ∈ ℝ^1000*×*750^ is used to model the EEG signal. Then, the dimension of X was reduced into *X* ∈ ℝ^*l×*750^ by selecting the top *l* eigenvectors with the highest eigenvalues. The value of *l* was chosen so that principal component 99% of the information content of the original *X* data set.

**Fig 3.**
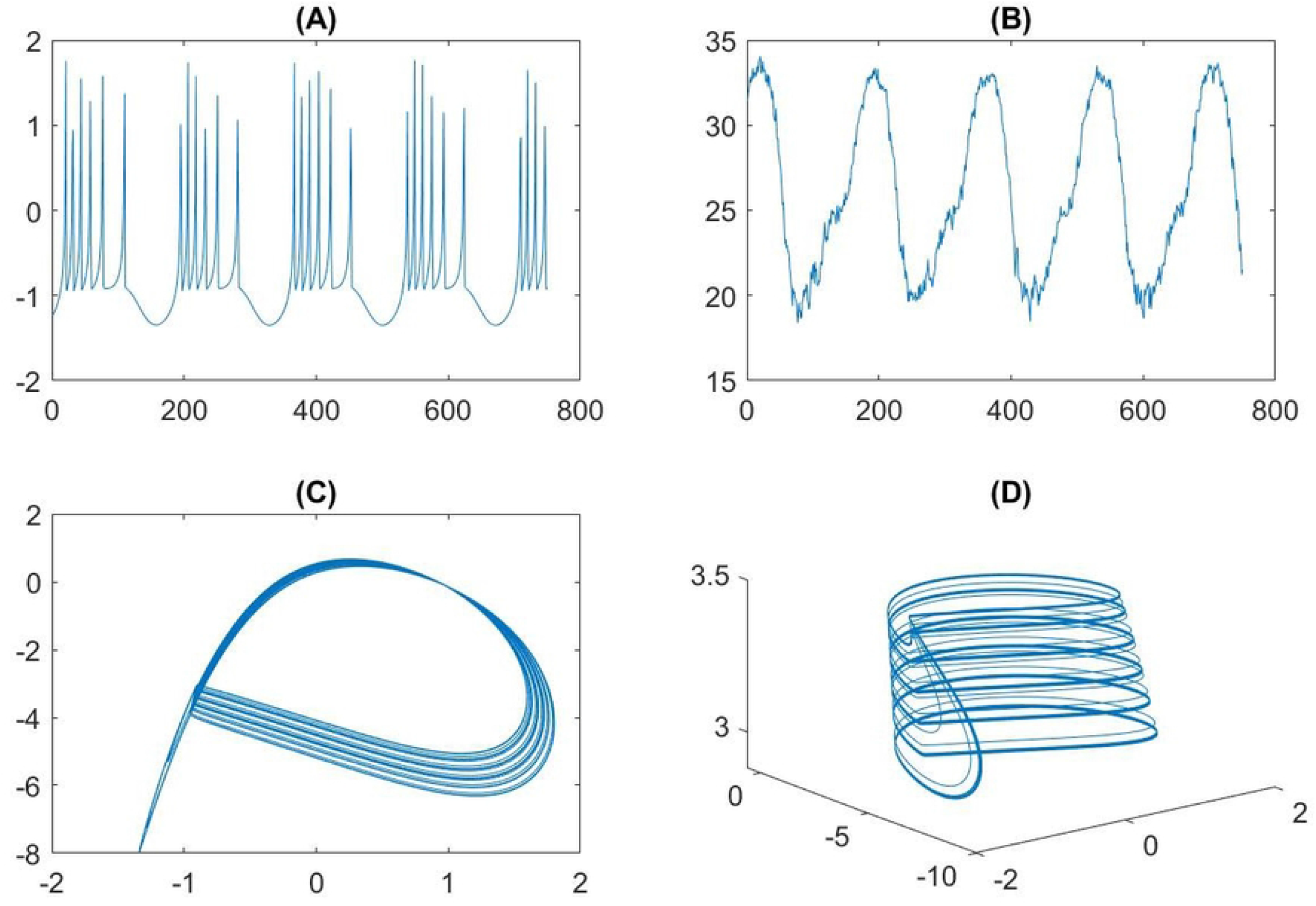
The activities of HR neuron. **(A)** action of membrane potential, **(B)** first principal component of membrane potential applying PCA, **(C)** and **(D)** a phase portrait.

The linear regression model was performed to extract the feature for each channel and sub-band frequency data. The weight coefficients were calculated by Eq. (3). Thus, we had a set of [*frequency × coefficient × channel*] arrays for every trials. Furthermore, the correlation between generated signals 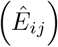 and original EEG was calculated as below:

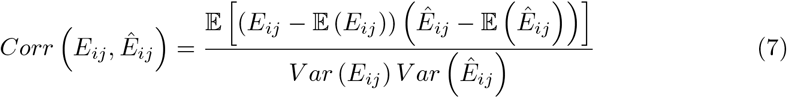

To evaluate the performance of proposed method for motor imagery classification, we conducted two experiments. In the first experiment, the session-to-session transfer classification, in which one session was for the training and one session was for the test, was conducted for each subject. Training and test sessions were recorded on different days for all subjects. Therefore, the session-to-session transfer classification is more challenging [51]. Whereas in the second experiment, we investigated across-subjects classification by using Leave-one-subject-out (LOSO) cross-validation, namely the data measured from eights subjects were used as the training set, while the data from the remaining one corresponded to the test set.

To demonstrate the efficiency of the proposed feature extraction method, accuracy and *Kappa* statistic defined as follows were used as the performance measure.

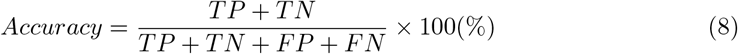

where TP indicates for true positive, meaning the correct classification as left-hand; TN indicates for true negative, meaning correct classification as right-hand; FP indicates for false positive, meaning incorrect classification as left-hand; and FN represents false negative, meaning incorrect classification as right-hand.

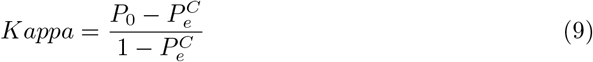

where *P*_0_ represents the probability of overall agreement between label assignment, classifier, and true process; and 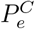 is defined by the chance agreement for all labels, i.e., the sum of the proportion of instances assigned to class multiplies in proportion to accurate labels of that specific class in the data set. Finally, to verify whether the performance difference between the proposed method and other methods is statistically significant, the two-side paired *t-test* has been conducted.

## Results

### Examine the results of extracted features

Figure 4 illustrates the distributions of correlation coefficients between original *E* and generated EEG signals *Ê* of subject 1. This result shows that the HR neuron model could fit the EEG signal with strong correlations (larger than 0.7). The highest correlation comes from the results in beta band (mean correlation coefficient was 0.89), while the smallest is found in theta band with mean values was 0.78. The mean values were 0.82 and 0.8 for alpha band and gamma band, respectively.

**Fig 4.**
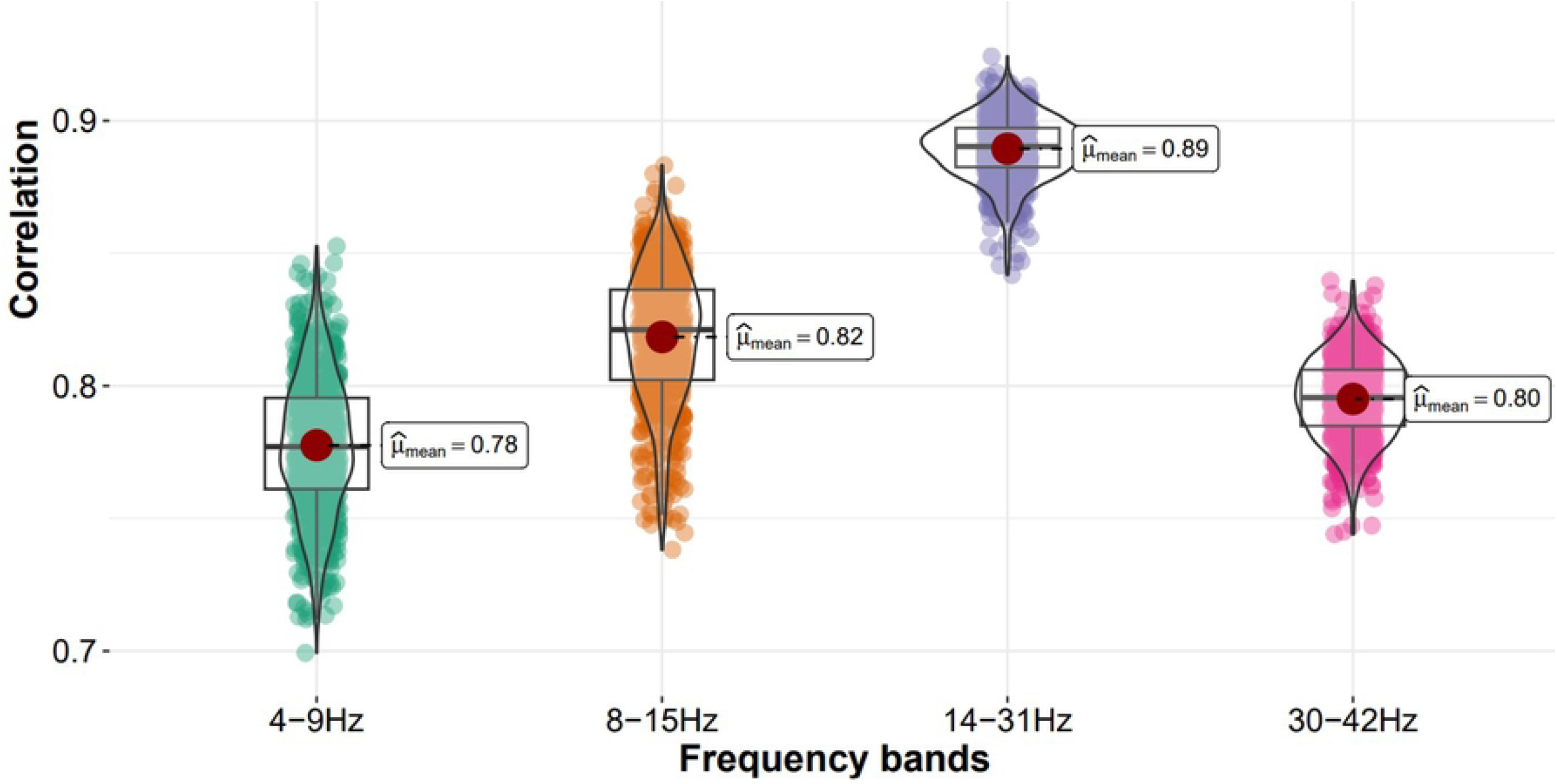
Box-plot of average correlation coefficient of C3, Cz, and C4 electrodes of subject 1 between actual EEG trials and generated EEG by HR neurons in four frequency bands. 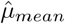 is the mean value of each frequency.

### Results on intra-subject classification

To evaluate the CAS-based feature for motor imagery classification, the intra-subject classification performance obtained by the proposed method was compared with FFT, STFT method and non-feature extraction works. The comparison is presented in Table 1 Considering the accuracy of 9 subjects, the proposed method yields the highest accuracy in 6 subjects.

**Table 1.** The accuracy and standard deviation (sd) in comparison on inter-subject with no feature extraction method [49], FFT [52], STFT [21]. Bold values represent the best result in each row.

| Subjects | No feature extraction |  | FFT | STFT | Proposed method |
| --- | --- | --- | --- | --- | --- |
|  | Shallow CNN | Deep CNN | CNN |  |  |
| 1 | 71.56 | 67.25 | 69.78 | 76.3 | **75.31** |
| 2 | 53.57 | 56.10 | 54.75 | 60.0 | **60.25** |
| 3 | 53.12 | 54.87 | 52.88 | **56.3** | 56.26 |
| 4 | 95.93 | 94.52 | 95.31 | 95.6 | **96.56** |
| 5 | 85.00 | 84.59 | 85.91 | 79.4 | **86.15** |
| 6 | 76.87 | 74.46 | 78.03 | 65.6 | **85.21** |
| 7 | **76.56** | 74.46 | 69.75 | 65.6 | 72.86 |
| 8 | 85.93 | **87.75** | 87.56 | 70.6 | 87.50 |
| 9 | 82.18 | 79.25 | 80.91 | 82.5 | **84.06** |
| Average ± sd | 75.64±14.41 | 75.10±13.58 | 74.99±14.52 | 72.4±12.31 | **78.23±13.28** |
| P – value | 0.02742 | 0.03254 | 0.00219 | 0.05254 |  |

In comparison, its accuracy in the other two subjects is very competitive (less than 4.13% and 0.25% compared with the highest ones). The results indicate that the proposed CAS-based feature extraction method is superior to other methods with 3.7% on the average level. The obtained p-value of two-sided paired *t-test* at the significance level of *α <* 0.5 The obtained p-values are given in the bottom row of Table 1. It can be seen that all the obtained p-values are less than 0.05, which suggests that CAS-based method significantly improves the classification accuracy in comparison with other method. Moreover, Zhu et al. utilized CSP method combined with CNN for the same dataset, achieving approximately 64% accuracy [14]. Compared with CSP method, our method yields a higher performance of 14.28% in classification.

Table 2 summarizes the average *kappa* values of the proposed method and non-feature extraction works. The kappa statistic has a value between 0 and 1, and the higher value indicates the better model consistency. Table 2 suggests that the proposed method performs better than non-feature extraction methods.

**Table 2.** The Kappa values in comparison on inter-subject with the CNN-based methods.

| Works | No feature extraction |  | Proposed method |
| --- | --- | --- | --- |
|  | Shallow CNN | Deep CNN | CNN |
| Kappa | 0.63 | 0.59 | 0.64 |

### Results on across-subjects classification

The data set contained EEG data of 9 subjects with 600 trials per subject. Using the LOSO protocol, the reduced features are partitioned into training data and test data. The data set has nine subjects, the validation process iterates nine times corresponding to 9-fold cross validation. For each iteration, the features data of one left-out subject is set as the testing data, and the features data of the remaining subjects are set as the training data.

In the test of across-subjects, Table 3 shows the accuracy of the proposed methods for 9-folds cross validation. Among 5400 trials, the CAS-based feature was classified using the CNN model. Specifically, study from [11] reported that DBN classification performance reached 84%, but the performance dropped to 70% when the model was trained in across-subjects setting. The HS-CNN reached 87.6% of intra-subject classification, but performance drops as 65.3% in across-subjects setting [10]. Our approach achieved 74.03% (±10.02%) mean accuracy (Table 4).

**Table 3.**
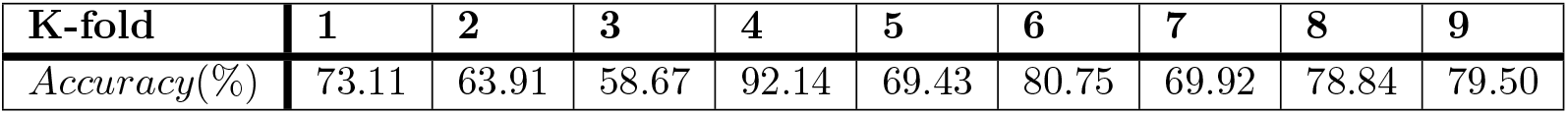
The accuracy on across-subjects with 9-fold validation of proposed method.

**Table 4.** The accuracy in comparison on cross-subject with state-of-the-art methods.

| Works | DBN | HS-CNN | Proposed |
| --- | --- | --- | --- |
| Accuracy(%) | 70 | 65.3 | 74.0 |

In comparison with other works, this study has two advantages. First, our study indicates that the CAS-based method is more robust and can extract distinguishable features from EEG signals more than FFT. The linear regression model uses a simple linear algebra formula and reduce the consumption of calculation, which is more beneficial for online BCI application. Second, our proposed method can improve the inconvenience of the parameter dependency such as time window, model order referred in [26].

The average processing time for the proposed methods was 15 ms per epoch on a PC with Intel® Core™ i7 4.20 GHz processor and 48 GB RAM. The code was developed on the Python Jupyter notebook.

## Conclusion

In this study, the EEG signals were reproduced using a linear combination of the states of dynamical neurons connected in a network where the HR neurons were operating in the CAS regime. We provide the evidence that MI-EEG data can be generated well using CAS model in four frequency bands, as demonstrated by the results from Figure 4.

This study used the CAS-based CNN approach to reconstruct the EEG signal as a minimal model of the brain. HR neurons weakly connected with spiking activities, and producing a mean field. We use the output from the CAS networks to model the EEG signal. In this sense, it has similarities to Reservoir Computer, a machine learning technique that does not train the synapses, but rather, the RC only optimizes the coefficients of the linear regression, but as the Reservoir is based on a nonlinear dynamics that can incorporate complex behaviour such as EEG signals. On the other hand, our integrated CAS and CNN method takes advantage of classification methods such as CNN to find hidden pattern. Different from applying CNN directly to the EEG signal, we apply CNN to the coefficients of the linear regression of the CAS model. But, since the regression is made on a reduced set of orthogonal components *l* which is smaller than the number of neurons and the number of points (sampled time series) of the original EEG data, classification can be performed on a small set of numbers (the coefficients of the regression) that have different significance (the first coefficient is a weight applied to the eigenvector that most contributes to the directions observed in the EEG experimental signal). This is different from applying CNN directly to the EEG signal, where the stochastic signals such as EEG signal contains 750 points, for example. Please note that each data point has equivalent significance. This allows our method to work even when the number of coefficients *l* (number of principal components) is very small, allowing for a classification that is less prone to errors and computational efficiency. The linear regression model has a certain limitation to capture the long-range time correlation [53]. The HR neurons network with CAS pattern are actually able to capture both short and long range temporal correlations.

Furthermore, the coefficients of the linear regression to model the EEG signals were used as feature vectors for classification. The performance of the proposed feature extraction method is compared with CNN-based methods for both intra-subject and across-subjects MI-EEG classification. Through the results of experiments, we first found that the CAS-based method successfully maintains important features of EEG signals, thereby improving the performance. Compared with non-feature extraction and FFT method, our proposed methods achieved superior classification performance for the intra-subject setting. Furthermore, our proposed method outperformed the classification accuracy with the state-of-the-art methods such as HS-CNN and DBN for across-subjects classification [10, 11].

In conclusion, this work demonstrates the superior performance and promising potential of the proposed EEG feature extraction (by the CAS model approach) and further classification method (by the CNN approach) using fewer assumptions. This article could pave the way for the practical implementation of an across-subject BCI.

In recent years, researchers have tried their best to design online BCI systems for commercial use [54]. The accuracy and non-stationarity characteristics of this signal are still a challenge for real-time BCI implementation. An online BCI system requires that the algorithm has a high performance [55–57]. The method proposed in this study could achieve superior classification performance for both intra-subject and across-subjects settings. Using the linear combination of the time series obtained from simulations of HR neural networks, our proposed method is independent from the model order, not the case for autoregressive methods. It also does not depend on the type of network being used to generate the simulated data, since this methodology is inspired by Reservoir Computing techniques that work regardless of the network topology, as long as neurons are sparsely connected. Moreover, the time-domain-based feature extraction approach allows a short-segmented data. Therefore, the proposed method would potentially contribute to the online BCI system design.

However, there are still several challenging issues that need to be addressed for future investigations. Firstly, the proposed method was compared with FFT features with one dataset. Further works on different features should be implemented to give reliable results. In addition, we used a simple CNN structure in this study. Several new architectures are worthwhile to explore to boost the performance further.

## Acknowledgments

The research was supported by JSPS KAKENHI (Grant numbers JP20H02111 and JP19H05727).

